# The ‘filtering’ metaphor revisited: competition and environment jointly structure invasibility and coexistence

**DOI:** 10.1101/356360

**Authors:** Rachel M. Germain, Margaret M. Mayfield, Benjamin Gilbert

**Affiliations:** Biodiversity Research Centre, University of British Columbia, Vancouver, British Columbia, Canada, V6T 1Z4; The University of Queensland, School of Biological Sciences, Brisbane, 4072 Queensland, Australia; Department of Ecology and Evolutionary Biology, University of Toronto, Toronto, Ontario, Canada, M5S 3B2

**Keywords:** annual plants, competition, fitness differences, niche differences, precipitation

## Abstract

‘Filtering’, or the reduction in species diversity that occurs because not all species can persist in all locations, is thought to unfold hierarchically, at large scales due to the environment and at small scales due to competition. However, the ecological effects of competition and the environment are not independent, and observational approaches preclude investigation into their interplay. We use a demographic approach with 30 plant species to experimentally test (i) the effect of competition on species persistence in two soil moisture environments, and (ii) the effect of environmental conditions on the mechanisms underlying competitive coexistence. We find that competitors cause differential persistence of species across environments even when these effects are lacking in the absence of competition, and that the traits that determine persistence depend on the competitive environment. Changing environmental conditions generated idiosyncratic effects on coexistence outcomes, increasing competitive exclusion of some species while promoting coexistence of others. Our results highlight the importance of considering environmental filtering in light of, rather than in isolation from, competition, and challenge community assembly models and approaches to projecting future species distributions.

## 1. Introduction

One of the most fundamental schematics shown in any introductory ecology class depicts the hierarchy of ecological filters thought to give rise to local communities from a regional species pool. The first filter generally depicted is the environment, which excludes any species without a suitably compatible phenotype, whereas the second filter represents competition among environmentally compatible species. This filtering metaphor has been applied to infer ecological process from a wide range of patterns, from the scale-dependence of biodiversity to the distribution of trait/phylogenetic relatedness in communities [1]. However, ‘filtering’, by definition, is the successful or failed persistence of species in a given environment, and is difficult if not impossible to detect without experimentation [2]. For instance, a species might be absent from a locality simply because it has not yet arrived [3] or be present but on an extinction trajectory that has not yet been realized [4]. Additionally, trait/phylogenetic patterns have been shown to lead to incorrect inferences about ecological mechanisms in the presence of competitors, which is the case in most environments [2,5], and studies often consider the environment and competition as alternative explanations for patterns, with little attention paid to their interaction.

Experiments are needed to understand how persistence and its demographic drivers are interactively structured by the competition and the environment [2]. In its broadest sense, filtering removes a subset of the potential community, and is quantifiable as population growth rates <1 when at low abundance (λ_*inv*_, known as ‘invasion growth rate’ [6]). By contrasting λ_*inv*_ of a given focal species among different environments, specifically in the presence and absence of competitors in different environments (λ_*inv[C+E]*_ and λ_*inv[E]*_, respectively), we can ask how competition changes predictions about which species can persist in a given environment. Broad-scale tests of the assumption that species are limited by environmental conditions, rather than community responses to those conditions, are rare [7] but needed to be understood to forecast the ecological consequences of environmental change (*e.g.*, species’ range limits [8]).

A complementary approach to testing persistence is to examine how the environment affects the mechanisms that underlie species coexistence (*i.e*., the mutual persistence of competing species [6]). Recent years have seen a considerable research effort put towards quantifying niche differences and fitness differences among species, which promote and preclude coexistence, respectively [9]. In doing so, ecologists are now much closer to resolving longstanding questions about how differences among species, such as functional traits [10], provenance [11], and evolutionary history [12], contribute to coexistence and the maintenance of diversity. However, it remains unknown how sensitive niche and fitness differences are to the environment, even in well-studied systems, as most experiments are conducted in single environments (but see [12,13]). Other research examines how environmental heterogeneity and dispersal allow spatial or temporal coexistence, underlain by species-specific environmental responses and dispersal (*e.g.*, storage effects [14], relative non-linearity). As a result, the scale-dependence of coexistence mechanisms are increasingly understood, but we lack empirical evidence of how sensitive local competitive interactions are to environmental context—these data are needed to understand the mechanistic interplay between competition and the environment.

To resolve the complex interplay between competition and environmental conditions, we couple a demographic approach [2,15] with trait data and an experimental manipulation of a key resource. We apply this approach to answer four questions: (1) How strong is environmental filtering *sensu stricto*, measured as the exclusion of species based on environmental conditions alone [2]? (2) How do interspecific competitors alter the effect of environmental conditions on persistence? (3) Do species’ traits explain differential responses to environmental conditions with and without competition? and (4) How sensitive are niche differences, fitness differences, and coexistence among competing species to the environment? We use annual plant communities from mediterranean-climate regions as a model system, contrasting two soil moisture regimes, ‘wet’ and ‘dry’, that represent differences between mesic (662 mm/year) and xeric (312 mm/year) sites across species’ ranges in California [16].

## 2. Materials and methods

Our experiment included 30 species grown for seven months in a research greenhouse at the University of Toronto (electronic supplementary material appendix S1). In brief, we grew two types of experimental communities: (1) species grown alone at low densities (7 plants/pot), such that individuals were not experiencing any interspecific competition and minimal intraspecific competition, and (2) species grown in pairs at high densities (70 plants/pot) at a range of relative frequencies to shift the strength of intraspecific and interspecific competition.

All experimental communities were replicated under wet and dry conditions, imposed by watering the wet treatment twice as frequently as the dry treatment. All seed was collected from senescing plants, enumerated, and used to estimate per capita seed production, from which we fit separate annual plant competition models (equation S1) for each species pair. Species’ traits were measured in separate experiments under identical growing conditions (electronic supplementary material appendix S1).

λ_*inv*_ estimates are species-specific, whereas niche and fitness differences are calculated for each pair of interacting species. λ_*inv[E]*_ is the finite rate of increase estimated via model fitting (λ_*i*_ in equation S1), whereas λ_*inv[c+E]*_ is the solution to 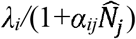,where 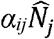 is the interspecific effect of species *j* on *i* (*α*_*ij*_) when species *j* is at its single-species equilibrium population size 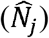. For each species pair, we used parameter estimates from the fitted competition model to solve for niche and fitness differences according to previous studies of annual plants [10,11]. Niche differences are derived from the relative strengths of interspecific and intraspecific competition (equation S2), whereas fitness differences are the product of species’ differences in fecundity and sensitivity to competition (equation S3). The joint values of niche and fitness differences can predict whether a species pair can coexist in a given environment (figure 2). We used linear mixed effects models to test differences in λ_*inv*_ and coexistence mechanisms among soil moisture environments (electronic supplementary materials, appendix S1). Note that our definition of ‘environment’ is inclusive of non-competitive species interactions, such as among plants and their soil microbes/pollinators.

## 3. Results and discussion

We found that filtering caused by the environment alone was dwarfed in comparison to the filtering effect of competition. Specifically, only a single species failed to persist (λ_*inv[E]*_<1) in the absence of competition compared to 12 per environment in the presence of competitors (pie charts in figures 1*a*,*b*). Soil moisture had no mean effect on λ_*inv*_ across species regardless of the competitor environment (figure 1*a*,*b*, table S2; *P*>0.70) despite large effects of soil moisture, competition, or both within species (figures 1*a*,*b* and S1)—this discrepancy occurred because equal numbers of species increased and decreased λ_*inv*_ in response to dry conditions.

**Figure 1.**
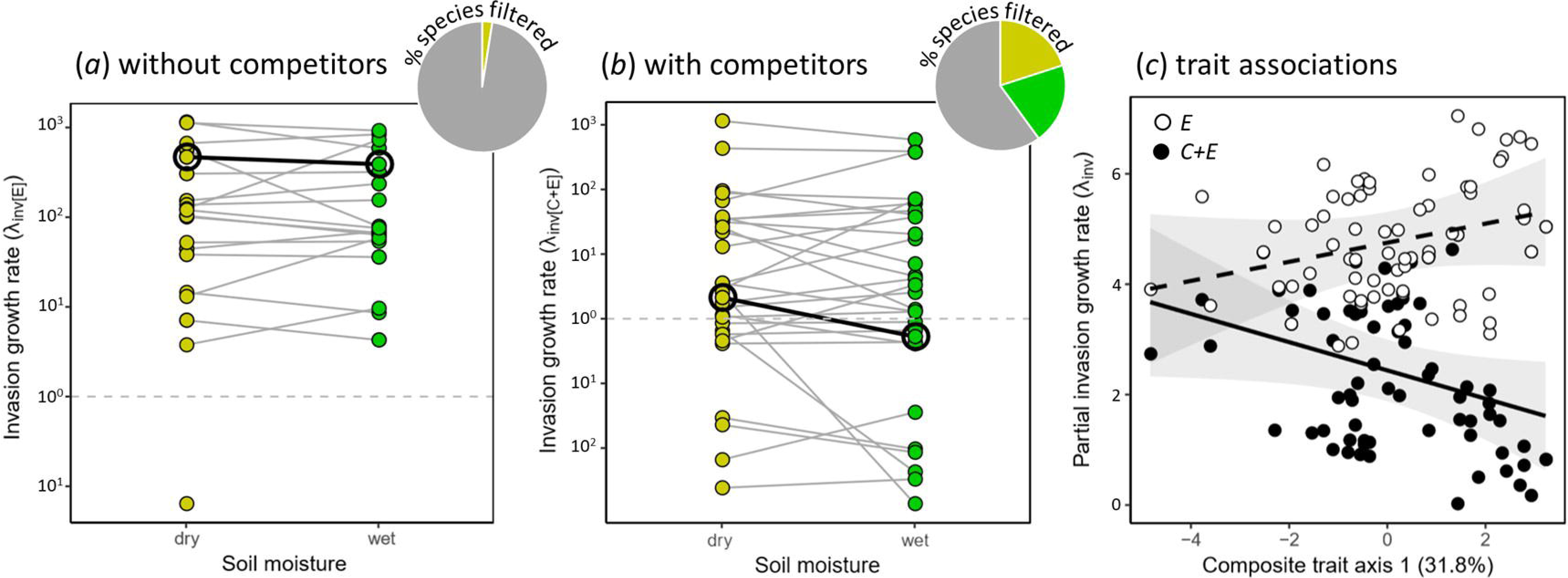
Invasion growth rate in the (*a*) absence (λ_*inv[E]*_) and (*b*) presence (λ_*inv[C+E]*_) of competitors in dry (yellow) and wet (green) soil moisture conditions, and (*c*) the underlying traits. (*a*,*b*) Points connected by a grey line show the same species in the different moisture treatments. Points ≥1 (dashed lines) are predicted to persist; *Lasthenia californica* is highlighted in bold. The subset of species pairs competed twice were not double-counted (table S1). (*c*) Points are λ_*inv[E]*_ and λ_*inv[C+E]*_ after partialling out variation explained by random effects, with fitted lines and 95% confidence bands. The composite trait is axis 1 from a multivariate analysis (figure S3).

Our results highlight two ways in which inferences of environmental filtering can be misled by the presence of competitors. First, for some species, the effect of soil moisture on
persistence only emerged in the presence of competitors. For example, *Lasthenia california* had high λ_*inv[E]*_ in both soil moisture environments, but in competition, was predicted to persist in dry conditions only (figure *1a,b).* Second, the traits that underlie species persistence (λ_*inv*_≥1) differed depending on whether competitors were present or absent (figure 1*c* and table S3; *p*=0.001). In observational studies, the strength of environmental filtering is typically inferred by tracking occupancy or measuring trait distributions in natural communities, which include competitors [1]. If our results had come from such an observational study, we would erroneously conclude that environmental filtering removes 40% of the species pool, specifically species with low biomass, shallow roots, and small seeds (biplot in figure S3). Previous research in species-poor communities suggests that inferring environmental filtering from trait patterns can mislead conclusions [5]; our experiment is the first to confirm this result using a robust demographic approach.

Niche and fitness differences responded strongly to the soil moisture environment for many species pairs (figure 2), shifting coexistence outcomes, despite lacking an average effect across species (table S4). Nine species pairs were predicted to coexist in each environment, but the identities of only six pairs were common to both environments. At first glance, these idiosyncratic responses were surprising—empirical experiments frequently predict that increasing resource supply rates reduces coexistence, specifically by decreasing niche dimensionality [17]. However, as our data suggests, this prediction entirely depends on where resource supply rates fall relative to species’ resource requirements and drawdown rates [18]. Additionally, theoretical predictions may not hold if phenotypic plasticity causes species’ resource requirements and drawdown to differ among environments (*e.g.*, converging root lengths [19]). Indeed, plasticity was high in our experiment for some species pairs (figure S3), but pairs exhibiting large competitive responses despite a lack of plasticity were also observed (*e.g., Lupinus* vs. *Trifolium* in figure S3), thus plasticity alone offers an insufficient explanation for observed environmental responses. In sum, the contrasting soil moisture environments tested here reflect those observed in the field and, although they have important consequences for coexistence, their effects are not generalizable across species.

**Figure 2.**
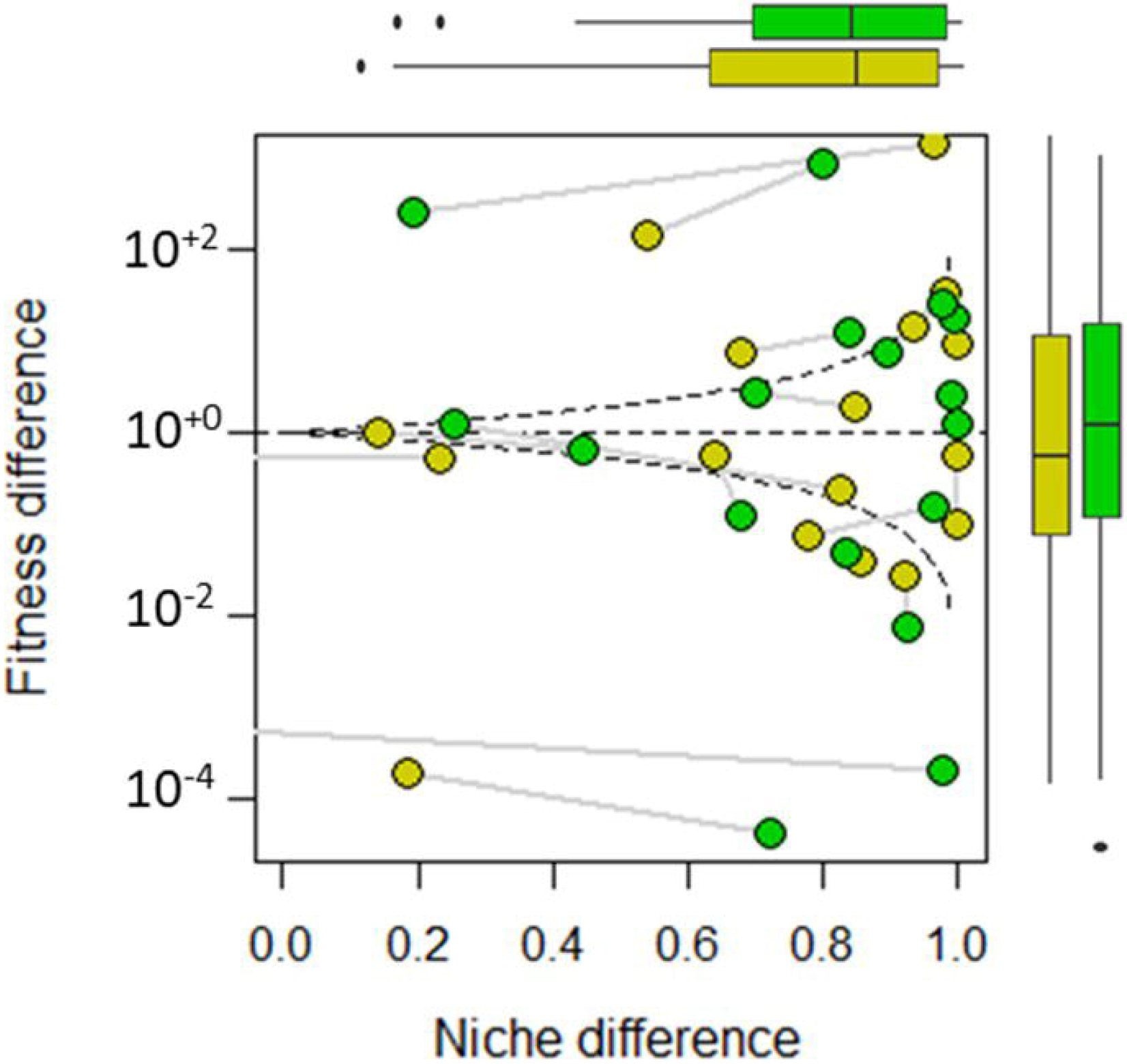
Joint responses of niche and fitness differences to wet (green) and dry (yellow) soil moisture treatments. Niche differences are maximized at 1, whereas fitness differences are a ratio (equation S3) and equal 10° when species have identical fitnesses. The curved dashed lines represent the boundary between coexistence and exclusion. Boxplots in the margins show distributions of niche and fitness differences among environments. Niche differences fell below 0 for four species pairs (figure S4), indicating positive frequency dependence.

The field of ecology is undergoing rapid conceptual revision, as classical ideas are deconstructed and rebuilt with greater theoretical support and empirical utility [15,18]. Our results speak to the need to critically re-evaluate the conceptual separation of environmental and competitive filters [15] and demonstrate the depth of mechanistic inference that can be drawn by disentangling their effects [2]. Future research opportunities include an examination of (i) a greater range of environmental gradients that species encounter in nature, (ii) multiple axes of resource limitation [17], and (iii) the mechanisms that underlie idiosyncratic environmental effects on coexistence mechanisms (*e.g.*, plasticity [19]). The integration of these approaches, including the results we report here, promise to lead to an understanding of how environmental conditions structure biodiversity and to more accurate forecasts of the impacts of global changes.

## Acknowledgments

We thank many undergraduate assistants for greenhouse assistance, notably Chris Blackford. Project feedback was provided by Angert, Williams, and Srivastava labs.

## Author contributions

R.M.G./B.G. designed the experiment, R.M.G. carried out the experiment and performed analyses, R.M.G./M.M.M./B.G. wrote the paper.

## Data accessibility

All data will be deposited on Dryad following manuscript acceptance.

